# Targeted nanopore resequencing and methylation analysis of LINE-1 retrotransposons

**DOI:** 10.1101/2022.06.25.497594

**Authors:** Arpita Sarkar, Sophie Lanciano, Gael Cristofari

**Author notes:** These authors contributed equally to the work (Arpita Sarkar, Sophie Lanciano).

## Abstract

Retrotransposition of LINE-1 (L1) elements represent a major source of insertional polymorphisms in mammals and their mutagenic activity is restricted by silencing mechanisms, such as DNA methylation. Despite a very high level of sequence identity between copies, their internal sequence contains small nucleotide polymorphisms (SNPs) that can alter their activity. Such internal SNPs can also appear in different alleles of a given L1 locus. Given their repetitive nature and relatively long size, short-read sequencing approaches have limited access to L1 internal sequence or DNA methylation state. Here we describe a targeted method to specifically sequence more than a hundred L1-containing loci in parallel and measure their DNA methylation levels using nanopore long-read sequencing. Each targeted locus is sequenced at high coverage (∼45X) with unambiguously mapped reads spanning the entire L1 element, as well as its flanking sequences over several kilobases. Our protocol, modified from the nanopore Cas9 targeted sequencing (nCATS) strategy, provides a full and haplotype-resolved L1 sequence and DNA methylation levels. It introduces a streamlined and multiplex approach to synthesize guide RNAs and a quantitative PCR (qPCR)-based quality check during library preparation for cost-effective L1 sequencing. More generally, this method can be applied to any type of transposable elements and organisms.

## 1. Introduction

Transposable elements (TEs) contribute to at least half of the human genome. Although the vast majority of these elements have lost their activity by accumulating mutations or truncations, hundreds of LINE-1 (L1) retrotransposon elements are still capable of mobilization in human genomes [1–4]. Their activity in germ cells or in the early embryo can lead to new inheritable genetic diseases. They can also replicate in somatic tissues, particularly in epithelial tumors, in which they can produce driver mutations [5–8]. A full-length and retrotransposition-competent L1 element is 6 kb in length. It starts with an internal promoter at its 5’ extremity, followed by two open reading frames (ORF1 and ORF2) encoding the retrotransposition machinery, and ends with a polyadenylation signal. L1 replication begins with its transcription as L1 RNA serves both as a template for reverse transcription and as an mRNA for the translation of L1 proteins. It can be repressed by epigenetic silencing pathways, which often implicate DNA methylation [9]. When a new L1 copy integrates into the genome, its sequence is virtually identical to that of its progenitor. Therefore, the most recently integrated L1 copies are almost identical to each other, and short sequencing reads derived from such sequences can often map at different genomic positions [10]. Over time, L1 elements accumulate mutations and truncations, which may or may not affect their ability to retrotranspose [11]. Thus, determining the internal sequence of L1 retrotransposons, as well as their methylation state, is essential for our understanding of the mutagenic potential and regulation of these elements.

Nanopore-based sequencing [12–14] has the potential to generate reads several kilobases long, which can easily span a whole L1 element and their genomic flanks and capture structural variations, haplotypes and single nucleotide polymorphisms linked to L1. Moreover, nanopore sequencing allows the direct identification of modified bases, such as 5-methylcytosine (5mC) without the need to treat DNA with bisulfite [15–18]. However, given the cost and the required throughput to achieve whole-genome nanopore sequencing (WGS) and methylation calling, targeted approaches can be preferable if a reduced number of loci are under investigation. Several strategies of targeted enrichment coupled with nanopore sequencing have been proposed, based on long-range PCR [19], capture by oligonucleotide probes or Cas9 [20, 21], or Cas9 digestion coupled to preparative pulse-field gel electrophoresis [22]. However, these methods are limited by their reduced yield, by difficulties in multiplexing or by the loss of information related to DNA modifications. In contrast, nanopore Cas9-targeted sequencing (nCATS) enables efficient enrichment and long-read sequencing of multiple loci in a single run of a MinION portable sequencer from Oxford Nanopore Technologies (ONT) [23]. First, purified high-molecular weight genomic DNA is extensively dephosphorylated. Then, the dephosphorylated DNA is digested *in vitro* with Cas9 ribonucleoprotein particles (RNPs) containing guide RNAs (gRNAs) targeting the borders of the regions of interest. Cas9 cleavage produces DNA ends with 5’-phosphate groups that enable the ligation of ONT sequencing adapters. The original nCATS protocol used synthetic gRNAs targeting up to 10 genomic regions varying from 12 kb to 24 kb [23]. Nevertheless, the cost of synthetic gRNA is an obstacle for extensive multiplexing. More recently, nCATS was adapted for the genome-wide localization of human-specific L1 elements (L1HS) using a unique gRNA targeting the internal sequence of these elements [24]. Although this approach has the potential to profile most TE elements present in the investigated genome, it focuses on TE-containing alleles. Here, we combine nCATS with a streamlined and cost-effective protocol to synthesize gRNAs in pools, to genotype more than 100 loci in parallel and provide the sequence and DNA methylation state of both empty and L1-containing alleles (**Figure 1**).

**Figure 1.**
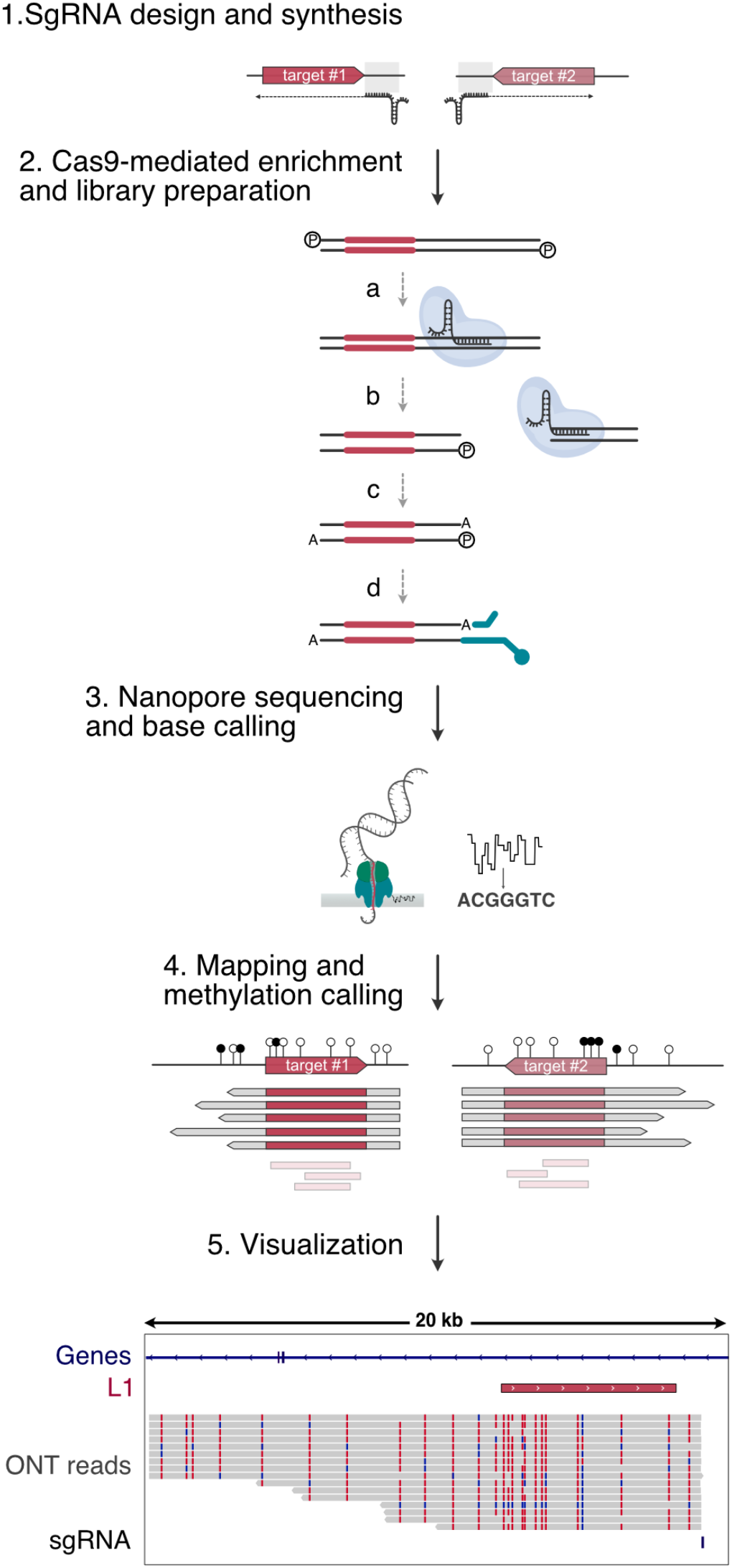
Overview of nanopore targeted sequencing of L1 loci. A schematic view of the main steps of needed for targeted nanopore resequencing and methylation analysis of L1 loci. **1**. An sgRNA is designed downstream of each targeted L1 (red solid arrows). All selected sgRNAs are synthesized *in vitro* in a single reaction. **2**. (a) Genomic DNA is extensively dephosphorylated and incubated with the pool of Cas9 ribonucleoprotein particles. (b) Cas9 cut leaves 5’-phosphate moieties at the cleaved DNA ends. For the sake of simplicity, the phosphate group attached to the Cas9-bound fragment is not shown. (c) An extra-A is added at the 3’ end of DNA molecules (A-tailing). (d) Oxford Nanopore Technology (ONT-sequencing adapters (green) are only added to the phosphorylated and A-tailed extremities. Cas9 bound to the other side of the cleavage site prevents ligation ensuring the directionality of the sequencing. **3**. The prepared library is loaded on a MinION flow cell and sequenced. **4**. ONT read are subsequently mapped to the reference genome and CpG methylation is called. Reads which map only to the internal sequence of L1 are discarded (light red). **5**. Read alignment, coverage and methylation levels can be visualized in a genome browser such as IGV. Tracks from top to bottom: Refseq, gene track with intron (blue line) and exons (blue boxes); L1, targeted L1 elements (red box); ONT reads, mapped nanopore reads (grey boxes). Vertical colored bars correspond to non-methylated CpG (blue) and methylated CpG (mCG, red). sgRNA, location of the sgRNA used to target this locus as a black bar.

Briefly, each locus of interest is probed by a single gRNA designed to bind downstream of and be antisense to the L1 element. All gRNAs are synthesized as a pool in a single *in vitro* reaction. In this setting, we obtain reads of varying length – but easily reaching several tens of kilobases -that preferentially extend from the region downstream of L1 to the region upstream of it, with an average coverage of 45X. Long-reads can be unambiguously mapped and used to accurately determine the DNA sequence and methylation profile of the whole L1 elements of interest, as well as those of their flanking sequences and matched empty alleles when the insertion is heterozygous. We also describe a quantitative PCR (qPCR)-based quality check to verify the efficiency of Cas9 digestion before ligating sequencing adapters.

## 2. Materials

All steps must be carried out with sterile nuclease-free plastics. Use:

1. DNA Zap (Life Technologies; to wipe bench-tops and pipettes)
2. Sterile low-binding 1.5 mL microtubes
3. Sterile low-binding filtered tips

### 2.1 Guide RNA synthesis and purification

1. Thermoblocks for 1.5 mL microtubes
2. Benchtop centrifuge for 1.5 mL microtubes with a rotor speed maximum 14,000*g*
3. Benchtop centrifuge with a swing bucket rotor and plate holders
4. Nuclease-free water
5. EnGen sgRNA synthesis kit (NEB)
6. Oligodeoxynucleotides (ODN) for sgRNA synthesis: positive control (LOU3161: ATTTGGACCTGCATCGTGCA) and Custom oligodeoxynucleotides (see section 3.1.2)
7. Monarch RNA Cleanup kit (NEB)
8. Qubit RNA Assay Kit (ThermoFisher)
9. Qubit fluorometer (ThermoFisher)
10. TBE-Urea gel 6%
11. TBE buffer

### 2.2 Genomic DNA extraction

1. Confluent T75 flask with the cell of interest for genomic DNA extraction
2. Monarch Genomic DNA Purification kit (NEB)
3. Qubit dsDNA HS Assay kit (ThermoFisher)
4. Vortex mixer
5. Agarose
6. TAE buffer
7. 0.07% Ethidium bromide solution
8. Electrophoresis apparatus-tank, gel tray, combs and power pack
9. DNA ladder
10. Nanodrop spectrophotometer

### 2.3 DNA Library Preparation

1. 0.2 mL PCR tubes
2. Thermocycler
3. CutSmart Buffer (NEB)
4. Nuclease-free TE buffer: 10 mM Tris-HCl, 0.1 mM EDTA, pH 8.0
5. Alt-R *S. pyogenes* HiFi Cas9 nuclease V3 (IDT)
6. Quick Calf Intestinal Phosphatase
7. Taq DNA polymerase
8. 100mM dATP solution, 100 mM
9. T4 DNA ligase from the NEBNext Quick Ligation Module (NEB)
10. MicroAmp Rapid 8-Tube Strip, 0.1mL (ThermoFisher) with MicroAmp 8-cap strip (ThermoFisher)
11. FastStart Universal SYBR Green Master (Rox) (Roche)
12. qPCR machine
13. Primers for qPCR: RASEF_forward (LOU3322: TCACAGGTTGCACACTGGAA), RASEF_reverse (LOU3323: AGCTCAGCCACTTTTCAGCT), Sox2_forward (LOU0695: CATGGGTTCGGTGGTCAAGT), Sox2_reverse (LOU0696: TGCTGATCATGTCCCGGAGGT)
14. AMPure XP beads (Beckman Coulter)
15. Magnetic rack for 1.5 mL microtubes
16. Ligation Sequencing kit (ONT)

### 2.4 Sequencing

1. Flow cell Priming Kit (ONT)
2. Flow cell FLO-MIN106 (pore version R9.4.1) (ONT)
3. Sequencer MinION Mk1B (ONT)
4. MinKNOW software (v 20.10.6) (ONT)
5. Mini-IT (v 10.42.0.1) (ONT)

### 2.5 Data analysis

1. Python 3 https://www.python.org/
2. Biopython (v 1.78) https://biopython.org
3. minimap2 (v 20.2) https://github.com/lh3/minimap2
4. nanopolish (v 0.13.2) https://nanopolish.readthedocs.io/en/latest/installation.html
5. Samtools (v 1.3) https://www.htslib.org/doc/samtools.html
6. Bedtools (v 2.30) https://bedtools.readthedocs.io/en/latest/
7. *reform* https://github.com/gencorefacility/reform
8. Integrative Genomics Viewer (IGV) software https://software.broadinstitute.org/software/igv/
9. Guppy (v 4.2.3) https://nanoporetech.com
10. NanoMethPhase https://github.com/vahidAK/NanoMethPhase.git
11. A multifasta file named `human.ref.genome.fa` with the reference human genome hg38 sequence (http://ftp.ensembl.org/pub/release-106/fasta/homo_sapiens/dna/Homo_sapiens.GRCh38.dna.toplevel.fa.gz)
12. A fasta file with the sequence of L1.3 (Genbank: L19088.1)

## 3. Methods

Typically, a sequencing run on a MinION flow cell consists of 5 steps: 1) planning the sequencing run, 2) prepare materials before library preparation, 3) library preparation, 4) sequencing, 5) data analysis.

### 3.1 Planning the sequencing run

About 2 weeks before the run, plan the experiment-select the target loci, design guide RNAs, order oligos to synthesize the guides. It is recommended that the flow cells and kits from ONT, as well as other consumables, be ordered well ahead of the actual run.

#### 3.1.1 Selection of target loci

As an example, we describe here an experiment designed to target 125 full-length L1-containing loci belonging to the youngest L1 family (L1HS), potentially polymorphic between individuals. The position of the target loci in the human reference genome (hg38) should be known.

#### 3.1.2 Design of guide RNAs

Cas9 is a site-specific DNA endonuclease that functions as a ribonucleoprotein particle (RNP). It finds its target -a 20 bp-sequence called *protospacer* - by base-pairing of its RNA moiety to the bottom strand of the protospacer. Cas9 binding to DNA also requires the presence immediately downstream of the protospacer of a short 3 bp motif, known as PAM (protospacer adjacent motif). The PAM sequence of *Streptococcus pyogenes* Cas9 (SpCas9) is 5’-NGG-3’ (top strand) [25, 26]. Absence of the PAM prevents binding of the Cas9 RNP to the DNA even if the RNA moiety perfectly matches the protospacer sequence [26].

Guide RNAs used for *in vivo* and *in vitro* experiments are 100 nt long artificial RNA molecules called “single guide RNAs” (sgRNAs), resulting from the fusion between a CRISPR RNA (crRNA) and a trans-activating-crRNA (tracrRNA) (**Figure 2**). The first 20 nucleotides at the 5’ end of the sgRNA are identical to the top strand of the protospacer and define Cas9 binding and cleavage specificities. This variable sequence (spacer) is followed by secondary structures derived from the 3’ end of the crRNA and from the tracrRNA, necessary for RNP formation and activation [26, 27]. When designing sgRNAs for SpCas9, the 20 nt variable sequence should be adapted to match the target site and should not be truncated. The remaining part remains constant.

**Figure 2.**
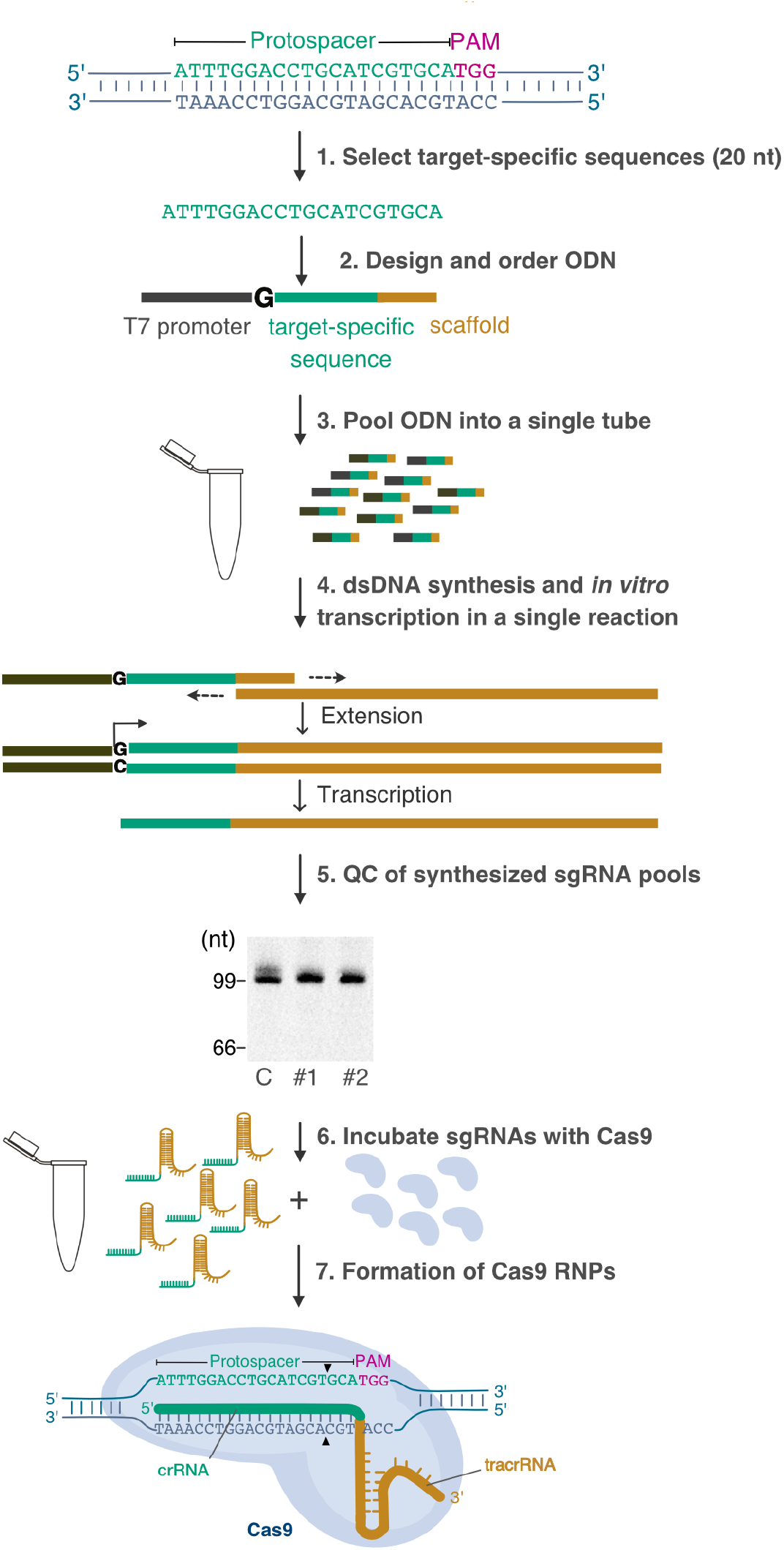
Preparation of Cas9 RNP pools. For each target locus, a Cas9 target sequence of 20 nucleotides (nt) is selected (1, see also **Figure 3**), and used to design an oligodeoxynucleotide (ODN) containing the T7 promoter sequence, the 20 nt Cas9 target sequence (without the PAM), and a 14 nt sequence complementary to the SpCas9 scaffold sequence (2). All ODNs are pooled in a single tube (3) and used to synthesize a pool of sgRNAs in a single-tube reaction using EnGen sgRNA Synthesis kit (4). After quality control by 6% denaturing gel electrophoresis in TBE-Urea (5), recombinant Cas9 is incubated with the sgRNA pool to form Cas9 RNP (6). Incubation of Cas9 RNPs with genomic DNA during subsequent library preparation leads to double-stranded DNA cleavage at the indicated position (black arrowheads) (7). C: CTRL ; #1: pool 1 ; #2 : pool 2.

Cas9 cuts the DNA duplex 3 bp upstream of the PAM sequence. Upon cleavage, it remains tightly bound to the protospacer-containing fragment and releases the PAM-containing one, which ends with a free 5’ phosphate that can be ligated to the nanopore sequencing adapter loaded with the DNA motor protein (**Figure 1**) [23, 25, 26]. Thus, nanopore sequencing begins at the PAM-containing end and continues in the opposite direction relative to the protospacer sequence. In summary, the direction of sequencing depends on the strand on which the PAM is located and therefore the sgRNA should be designed accordingly.

The following steps describe how to design an sgRNA in order to sequence a polymorphic L1 locus in the human genome, taking advantage of precomputed SpCas9 sgRNA target prediction and scoring by the CRISPOR tool [28] available in the ‘*CRISPR Targets*’ track of the UCSC Genome Browser (*see* **Note 1**, and **Figure 3**). For every sgRNA targeting the human genome, CRISPOR predicts:

**Figure 3.**
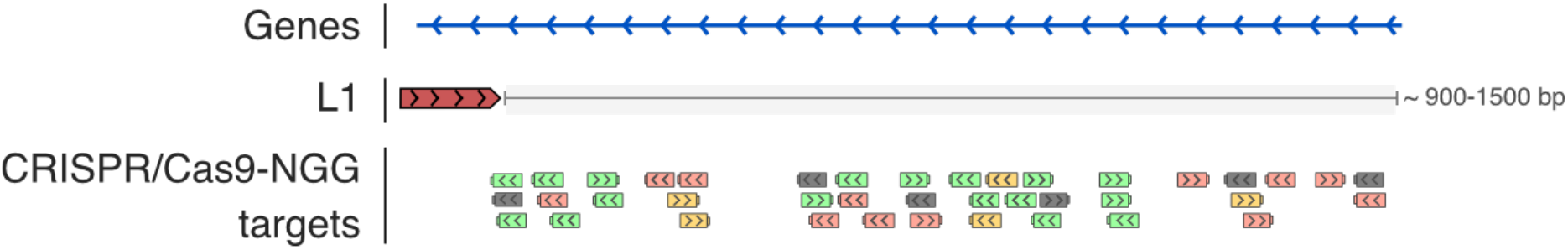
Selection of sgRNA downstream of L1 loci of interest. A UCSC Genome Browser view of the region downstream of an L1 element of interest (red solid arrow), showing CRISPR/Cas9-NGG target track. Potential CRISPR/Cas9 targets are shown as boxes colored according to their predicted efficiencies: low (pink), medium (yellow) or high (green). Non-unique targets are colored in dark grey. Each target is 20 nt long and can be found on the plus or minus strand (arrowheads inside boxes) with adjacent PAM (knob). We recommend selecting green target sequences with a minimum MIT score of 55 for and a minimum MM score of 35. To sequence L1 from the downstream region, an antisense sgRNA relative to L1 should be chosen. Note that L1 can also be sequenced from its upstream flank, choosing an upstream sgRNA. In this case, the sgRNA should be chosen in the sense orientation relative to L1.

a. a score of specificity, called the MIT specificity score [29]: the higher the score value (0-100), the higher the specificity of the sgRNA (and therefore the lower risk of off-targets).
b. a score of efficiency, called Moreno-Mateos Efficiency score or MM score [30]: the higher the score (in percentile), the greater the efficiency of the sgRNA.

We use the EnGen sgRNA Synthesis kit to synthesize sgRNAs. Synthesis reactions contain a single-stranded oligodeoxynucleotide (ODN) with the SpCas9 scaffold sequence, provided in the kit, and are supplemented with a custom ODN comprising the T7 promoter sequence, the target-specific sequence of the desired sgRNA (20 nt spacer obtained above), and a sequence complementary to the scaffold ODN (14 nt) (**Figure 2**). In a first step the two ODNs are annealed and extended by a DNA polymerase to form a complete double-stranded DNA template used, in a second step, for *in vitro* transcription by T7 RNA polymerase. Although this approach was designed to synthesize one sgRNA per reaction, we adapted it to synthesize a pool of 125 guides in a single reaction (*see* **Note 2**).

1. Go to the UCSC Genome browser choosing hg38 as a reference genome.
2. Display the *CRISPR Targets* track available in the *Genes and Gene Predictions* section, and the *RepeatMasker* track in the *Repeats* section.
3. Navigate to the coordinates of the L1 copy of interest (*e*.*g*. chr9:83049540-83055571). This TE appears in the *Repeats* track as a horizontal filled black box.
4. Zoom in to the target locus until the color-coded gRNA targets become visible, focusing to the region downstream of the TE, ∼900 to 1,500 bp from its 3’ end (*see* **Note 3**).
5. Select the sgRNA target with the highest scores, and with at least 55 and 35, for the MIT and MM scores, respectively, preferentially outside of other repeats. If high-scoring sgRNA targets cannot be found in this immediate flanking region, extend the search to a broader region.
6. Choose the direction of the guide (plus or minus strand). The L1 copy chosen as example is in the antisense orientation. As we would like the long reads to cover the entire L1 sequence together with its 5’ genomic flank, we choose a plus (+) strand sgRNA.
7. Copy the 20 nt spacer sequence, but not the PAM sequence, from the selected sgRNA, and store it in a data sheet with an appropriate name.
8. For multiple targets, repeat the operation or automate the process (*see* **Note 4**).
9. Once the sgRNAs have been selected, design DNA oligonucleotides for *in vitro* transcription of the sgRNAs. Starting from the 20 nt spacer (target) sequences selected at step 7, add one “G” at the 5’ end if this sequence starts with a different nucleotide, as a G is essential for transcription initiation by T7 RNA polymerase.
10. Add the T7 promoter sequence to the 5’ end: 5’-TTCTAATACGACTCACTATA-3’.
11. Add the 14 nt scaffold overlapping sequence to the 3’ end: 5’-GTTTTAGAGCTAGA-3’
12. The resulting ODN is 54 or 55 nt long and ready to be ordered.
13. Repeat the process for each of the 124 L1 target loci.
14. Add an additional ODN as positive control (LOU3161) to the same plate (*see* **Note 5**).
15. Order ODNs in a 96-well plate format (scale of synthesis: 25 nmoles per oligo, dried, standard-desalted).

### 3.2 Prepare materials before library preparation

#### 3.2.1 sgRNA synthesis and purification

The principle of sgRNA synthesis is depicted in **Figure 2**.

1. Spin the ODN plates briefly in a centrifuge with a swinging bucket rotor and plate holders and open carefully under a PCR hood.
2. Resuspend each ODN in nuclease-free water or TE pH 8.0 (IDT) at 200 µM (approximately 95 µL each). Pipette up and down each well carefully and incubate the plates at room temperature for 10 min to allow the ODNs to dissolve completely. Transfer the reconstituted ODNs to pre-labelled tubes.
3. Transfer 2 µL of each ODN at the stock concentration (200 µM) into a new 1.5 mL microtube and adjust the total ODN concentration to 50 µM by adding the required volume of water (750 µL of water for a pool of 125 distinct ODNs).
4. Transfer 20 µL of the ODN pool at 50 µM into 980 µL of water, to obtain a working ODN pool at 1 µM.
5. Thaw the EnGen 2X sgRNA Reaction Mix, *S. pyogenes* and 0.1 M DTT. Warm all the reagents except the enzyme mix at 37 °C for 10-15 min, mix and spin down in a microcentrifuge (*see* **Note 6**). Assemble the reactions at room temperature.
6. For *in vitro* transcription of the sgRNA pool, prepare duplicate 0.2 mL PCR tubes, with the following reagents in the order listed:

7. Mix each tube thoroughly and spin down in a microcentrifuge. Incubate at 37 °C for 2 h. Then keep on ice.
8. Bring the volume of each reaction to 50 µL by adding 30 µL of nuclease-free water, and add 2 µL of DNase I (supplied in the kit). Mix thoroughly and incubate at 37 °C for 15 min.
9. Proceed immediately to column purification.
10. Purify the synthesized sgRNAs with the Monarch RNA Cleanup kit. The protocol below purifies RNAs ≥ 25 nt. The columns supplied have a maximum capacity of 50 µg of RNA. To obtain highly concentrated RNA, the two duplicate reactions with the sgRNA pools are loaded to the same column.
11. Add 100 µL of RNA Cleanup Binding Buffer to each of the *in vitro* transcription reactions.
12. Add 150 µL of ethanol (≥ 95%) to each tube and pipet up and down to mix.
13. Prepare a single column with its collection tube.
14. Load one of the sgRNA pool reactions to the top of the column.
15. Spin at 16,000 x *g* for 1 min at room temperature and discard the flow-through.
16. Load the second sgRNA pool to the top of the same column.
17. Spin at 16,000 x *g* for 1 min at room temperature and discard the flow-through.
18. Wash the column with 500 µL of RNA Cleanup Wash Buffer.
19. Spin at 16,000 x *g* for 1 min at room temperature and discard the flow-through.
20. Repeat steps 18 and 19.
21. Spin again the column at 16,000 x *g* for 1 min at room temperature to eliminate any traces of ethanol and salts.
22. Transfer the column to a new RNase-free 1.5 mL microtube.
23. Load 50 µL of nuclease-free water to the top of the column and incubate for 2 min at room temperature.
24. Spin at 16,000 x *g* for 1 min at room temperature to elute the RNA.
25. Save 2 µL of the purified RNA for quantification and quality check, and prepare 5 µL-aliquots with the rest. Store them at -80 °C.
26. Quantify the sgRNA preparation with the Qubit RNA Assay kit and fluorometer according to manufacturer’s instructions. The expected yield ranges between 14 and 18 µg of RNA (*see* **Note 7**).
27. Perform a quality check by running 100 ng of the synthesized sgRNA pool on a denaturing 6% Novex TBE-Urea gel. The RNA should look like a compact band around 100 nt; any smear below this band indicates RNA degradation (**Figure 2**). Occasionally a slight smear may appear above the band indicating incomplete denaturation of sgRNA secondary structures.

|  |  |
| --- | --- |
| Nuclease-free water | 2 $\mu\text{L}$ |
| Engen 2X sgRNA reaction Mix, *S. pyogenes* | 10 $\mu\text{L}$ |
| ODN pool (1 $\mu\text{M}$) | 5 $\mu\text{L}$ |
| DTT (0.1 M) | 1 $\mu\text{L}$ |
| EnGen sgRNA Enzyme Mix | 2 $\mu\text{L}$ |
| **Total** | **20 $\mu\text{L}$** |

#### 3.2.2 Genomic DNA extraction

The biological samples should be collected and stored in conditions that do not alter high molecular weight genomic DNA. Whenever possible, working with fresh material is often preferred. We used the pellet of 5 × 10^6^ cultured cells as starting material for the Monarch Genomic DNA Purification kit:

1. Add 1 μl Proteinase K and 3 μl RNase A to the resuspended cell pellet and mix by immediately vortexing briefly. From this step onwards, the tube is kept at room temperature on the bench.
2. Add 100 µL of Cell Lysis Buffer and vortex well immediately.
3. Incubate the tube in a thermal mixer at 56 °C and agitate vigorously (∼1400 rpm) for 5 min. Simultaneously, heat nuclease-free water to 60 °C in another heat block.
4. Add 400 µL of genomic DNA Binding Buffer and vortex well for 5-10 s to mix the sample thoroughly.
5. With a pair of sterile scissors, cut the ends of 1 mL filtered tips to widen their opening. Use these wide-mouth tips to gently transfer the lysate into a genomic DNA Purification Column.
6. Spin the column: at first for 3 min at 1,000 x *g* to bind genomic DNA, then for 1 min at maximum speed (> 12,000 x *g*). Discard the flow-through and the collection tube.
7. Transfer the column to a new collection tube and add 500 μL of genomic DNA Wash Buffer on top of the column. Mix the closed column by inverting it very gently several times, so that the wash buffer touches the cap. Spin at 12,000 x *g* for 1 min and discard the flow-through.
8. Add 500 μL of genomic DNA Wash Buffer again, spin at 12,000 x *g* for 1 min and discard the flow-through.
9. Transfer the column to a fresh 1.5 mL microtube. Add 36 µL of the pre-heated water on the top of the column, and incubate at room temperature for 1 min. Spin at 12,000 x *g* for 1 min to elute the genomic DNA into the 1.5 mL microtube.
10. Save 3 µL of DNA for quantification and quality check and store the rest at 4 °C.

#### 3.2.3 Quality check and quantification of genomic DNA

1. Measure genomic DNA concentration by fluorimetry using the Qubit DNA assay and fluorometer according to manufacturer’s instructions. Starting from 5×10^6^ cells, the yield is ∼ 10 µg of genomic DNA. It is convenient to have concentrated DNA for the following steps (≥200 ng/µL, i.e. 5 µg / 24 µL).
2. Verify genomic DNA purity using 1 µL by measuring absorbance in a Nanodrop spectrophotometer (OD 260/280 nm = 1.8, and OD 260/230 nm = 2.0–2.2).
3. Resolve 100 ng of genomic DNA with a high-range DNA ladder (such as Eurogentec SmartLadder 200 bp-10 kb) by 0.8% agarose gel electrophoresis in 0.5X TAE. Typically, high molecular weight genomic DNA appears as a compact band higher than 10 kb. A smearing pattern below this size indicates breaks in the DNA and will result in a shorter read length.

### 3.3 Library preparation

Prepare wide-mouthed 1 mL and 200 µL filtered tips by cutting their end with a sterile scissors, and use them to gently pipette genomic DNA to avoid fragmentation. Each library is prepared from a single genomic DNA sample and independently sequenced in a single run.

#### 3.3.1 Assembly of Cas9 ribonucleoprotein particles (RNPs)

1. Set up the following reaction in a 0.2 mL PCR tube

2. Mix well by flicking the PCR tube and incubate in the thermal cycler at 25 °C for 30 min. Keep at 4 °C until further use.

|  |  |
| --- | --- |
| CutSmart Buffer 10X | 3 µL |
| Cas9 HiFi (62 µM) | 1 µL (final 2 µM) |
| sgRNA (pool of 125 guides) | up to 26 µL (final 2 µM) |
| Nuclease-free water | complete to 30 µL |
| **Total** | **30 µL** |

#### 3.3.2 Dephosphorylation of the free 5’ ends of genomic DNA

1. Set up the dephosphorylation reaction during incubation time of RNP assembly in a 0.2 mL PCR tube:

2. Mix gently by flicking or tapping the PCR tube (do not pipet up and down)
3. Spin down in a microcentrifuge
4. Incubate the reaction in a thermal cycler at 37 °C for 10 min, then at 80 °C for 2 min (to inactivate the CIP), and finally keep at 20 °C until use.

|  |  |
| --- | --- |
| High molecular weight genomic DNA (use wide-mouth tips) | up to 24 µL |
| CutSmart Buffer 10X | 3 µL |
| Quick CIP (5 U/µL) | 3 µL |
| Nuclease-free water | complete to 30 µL |
| **Total** | **30 µL** |

#### 3.3.3 Cas9-mediated digestion and dA-tailing of the target DNA region

1. Prepare a dATP working solution (10 mM final) by diluting 1 µL of the dATP stock solution (100 mM) into 9 µL of nuclease-free water and keep on ice.
2. Add directly to the previous reaction (from step 3.3.2) the following reagents:

3. Gently mix the PCR tube by flicking or tapping, and spin down in a microcentrifuge.
4. Save a 2 µL aliquot and keep on ice (as an undigested control for the qPCR assay).
5. Incubate the PCR tube in a thermal cycler with the following program: 1h at 37 °C, 5 min at 72 °C, on hold at 4 °C.
6. Save a 2 µL aliquot of the digested genomic DNA (for the qPCR assay), and reserve the rest at 4 °C until further use.

|  |  |
| --- | --- |
| Dephosphorylation reaction mix with the genomic DNA (from step 3.3.2) | 30 µL |
| Cas9 RNP formed with the pool of sgRNAs (from step 3.3.1) | 10 µL |
| dATP (10 mM working solution) | 1 µL |
| Taq Polymerase (5 U/µL) | 1 µL |
| **Total** | **42 µL** |

#### 3.3.4 Quality check of cleaved DNA by qPCR assay

Before ligating sequencing adapters, we recommend determining the efficiency of Cas9 cleavage. As a global proxy for sgRNA synthesis, RNP assembly and Cas9-mediated cleavage, we measure the copy number of the *RASEF* locus targeted by the control sgRNA LOU3161 within the pooled sgRNAs, before and after Cas9 digestion (*see* **Note 5**). This is achieved by qPCR relative quantification using a SYBR Green reaction mix. Successful cleavage by Cas9 results in a 15-fold decrease in *RASEF* copy number in the digested sample as compared to the undigested sample. We assume that all the other targets are similarly digested with the efficiencies predicted from sgRNA design. Failure of cleavage indicates poor performance and should be fixed before proceeding further. The qPCR assay amplifies two loci: 1) *RASEF* to measure cleavage, and 2) *SOX2* to normalize for genomic DNA quantity.

Under a PCR hood:

1. Dilute 60 times the genomic DNA samples saved before and after Cas9 digestion by adding 118 µL of nuclease-free water to each of the 2 µL aliquots.
2. Prepare two master mixes, one for each locus, with duplicate reactions, and a water control:

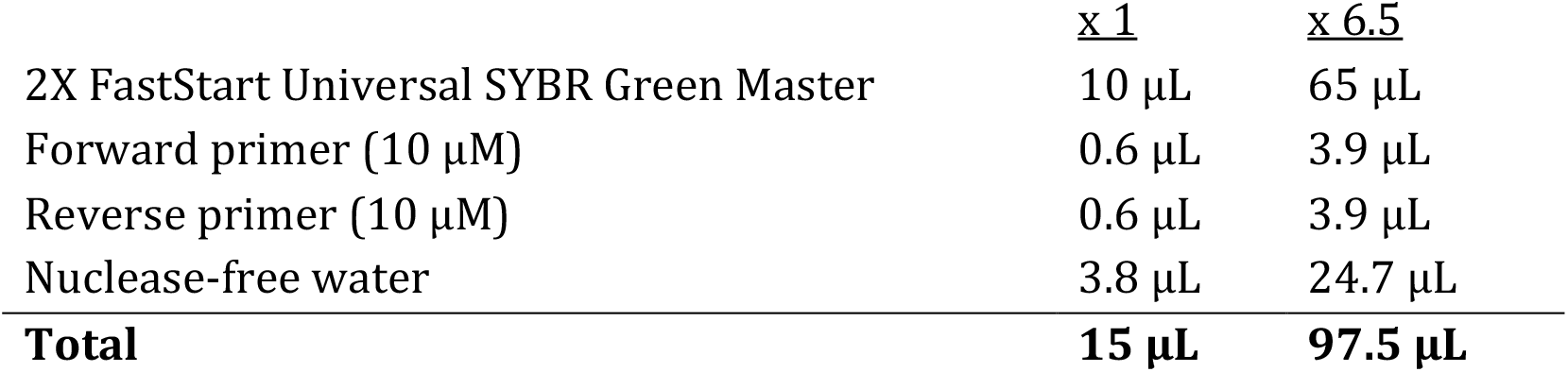
3. Set up the qPCR reactions in an 8-tube qPCR strip (**Figure 4a**).

**Figure 4.**
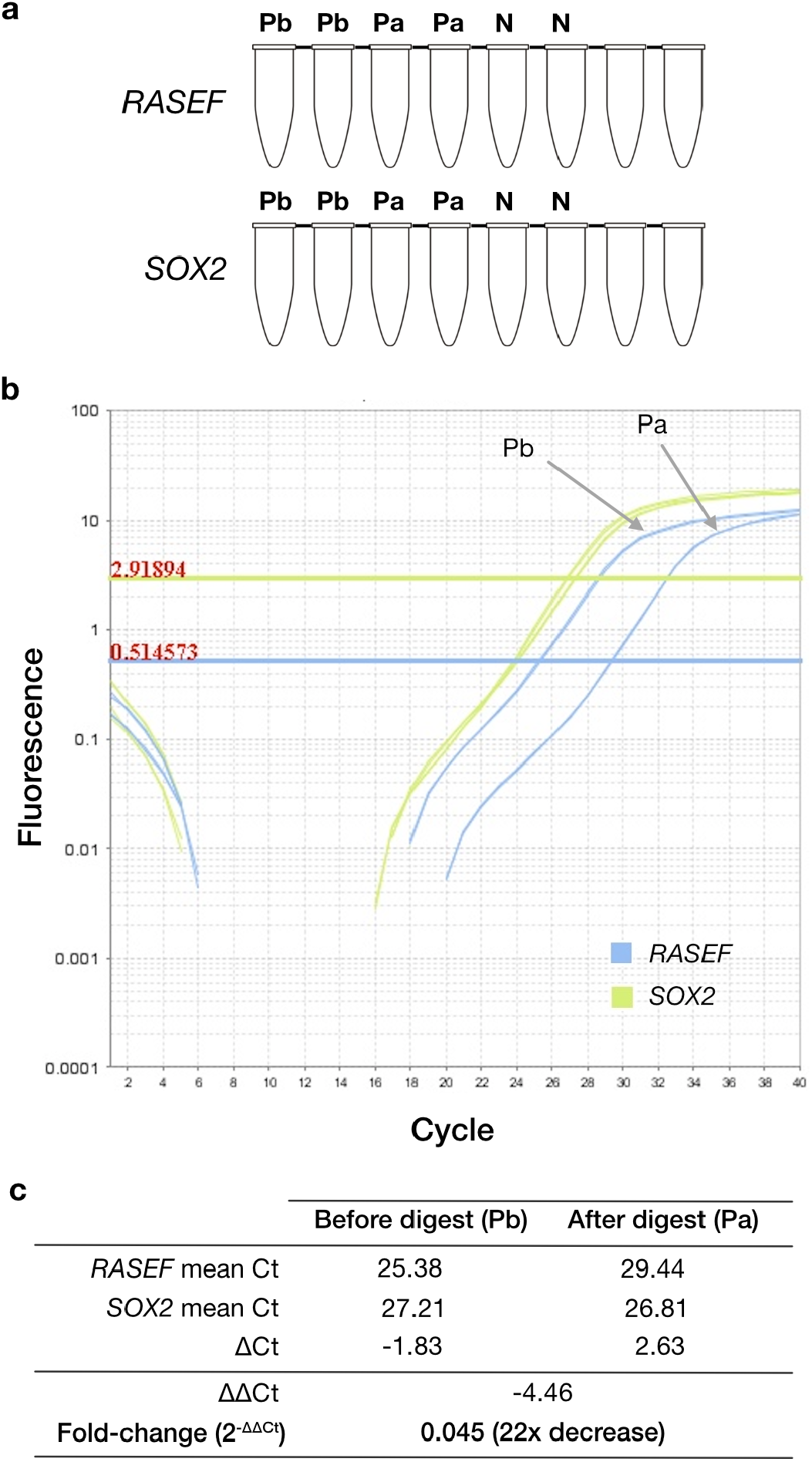
qPCR design and expected results. **(a)** Organization of the 8-tube strips for the *RASEF* and *SOX2* primer mixes. Pb: DNA sample before digestion with the pool of Cas9 RNP. Pa: DNA sample after digestion with the pool of Cas9 RNP. N: No template control. Each sample is loaded in duplicates. **(b)** Example of amplification curves generated by the StepOne software. Raw Ct values (averaged over duplicates) are shown on the top. Threshold settings for each primer mix were determined automatically by the software and displayed as horizontal lines. **(c)** Calculation of the fold change of intact DNA from the raw Ct values.
4. Dispense 15 µL of master mix, and 5 µL of diluted DNA or water per tube. Close the strip and mix by tapping. Spin briefly in a micro-centrifuge and insert the strip into the qPCR thermal block.
5. Open the StepOne software (for Applied Biosystems qPCR machines). Create a new experiment and enter program details such as ‘SYBR Green’ and ‘Relative quantification-Comparative Ct’. On the plate-layout, label each sample by its name, the type and name of the target locus.
6. Start the qPCR program: 10 min at 95 °C, 40 x [15 s at 95 °C, 1 min at 60 °C].
7. At the end of the run, click on ’Analyze’. The StepOne software generates amplification curves and tabular data for values of Ct across all tubes (**Figure 4b**).
8. Usually the ∆∆Ct value when comparing undigested and digested samples is ≥4.3, indicating successful cleavage by Cas9 (intact DNA represents less than 5% of the starting material after digestion, **Figure 4c**).

#### 3.3.5 Ligation of Sequencing adapters

In this step, sequencing adapters are specifically ligated to the DNA ends generated by Cas9 cleavage.

1. Place AMPure XP beads at room temperature.
2. Thaw the Ligation Buffer LNB at room temperature. Mix this viscous buffer by pipetting and spin down in a microcentrifuge. Place on ice.
3. Thaw the Adapter mix AMX-F and place on ice. Flick the tube to mix and spin down in a microcentrifuge. Place on ice.
4. Very gently, transfer the A-tailed DNA (38 µL from step 3.3.3) into a 1.5 mL microtube using wide-mouth tips.
5. Prepare the ligation reaction in another 1.5 mL microtube by adding the following reagents in the order stated:

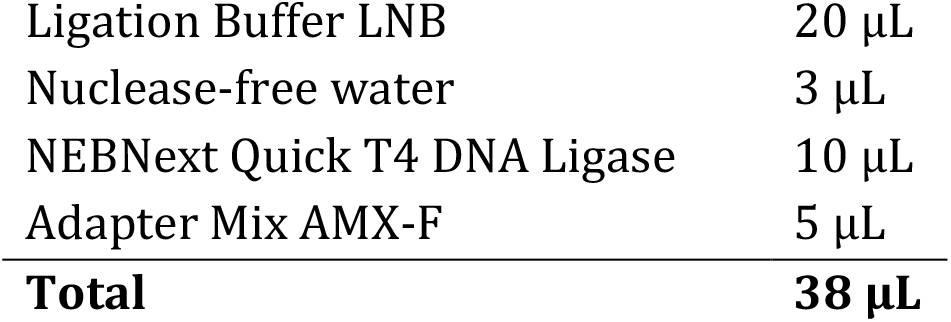 Mix thoroughly by pipetting up and down several times.
6. Add 20 µL of the above ligation mix to the DNA sample and mix gently by tapping the tube. Then add the rest of the ligation mix to the sample and mix gently again. The 2-step incorporation helps mixing despite the high viscosity. Spin down in a microcentrifuge.
7. Incubate the reaction for 10 min at 20°C.

#### 3.3.6 AMPure XP bead purification of the DNA library

1. Vortex the bottle of AMPure XP beads (already brought to room temperature) to resuspend the beads.
2. Thaw the Long Fragment Buffer LFB and Elution Buffer EB at room temperature.
3. Adjust the volume of the ligation reaction mix containing the genomic DNA to ∼160 µL by adding 80 µL of TE (pH 8). Mix gently by flicking or tapping the tube.
4. Add 48 µL of AMPure bead suspension to the sample to obtain a beads:DNA ratio of 0.3:1. Mix gently by flicking or tapping the tube.
5. Incubate for 10 min at room temperature.
6. Spin down gently in a microcentrifuge if necessary, and transfer the tube to a magnetic rack. Wait ∼5 min to pellet the beads with the magnetic field.
7. Carefully remove the clear supernatant with a pipette.
8. Add 250 µL of LFB buffer. Washing with LFB eliminates DNA fragments shorter than 3 kb. Flick or tap the tube gently to resuspend the beads, then place back on the magnetic rack for 5 min.
9. Carefully remove the clear supernatant with a pipette.
10. Repeat steps 8 and 9.
11. Do not remove the tube from the magnetic rack. With a 10 µL pipette, remove the last drops of supernatant and let the pellet to dry for 30 s while the tube is still on the rack. Do not dry more than 30 s, otherwise the pellet may over dry and crack. The pellet should look dry but shiny.
12. Remove the tube from the magnetic rack and add 13 µL of EB buffer to resuspend the pellet. Mix gently by flicking or tapping and incubate at room temperature for 30 min.
13. Place the tube back in the magnet rack, pellet the beads for 5 min, ensuring that the eluate is clear.
14. Pipette the eluate from the tube into a clean 1.5 mL microtube.
15. Take 1 µL of the eluted library for quantification and store the rest at 4 °C.
16. Quantify the DNA library with a Qubit fluorometer and Qubit DNA assay. It is recommended to load between 20 and 50 fmol of DNA library on a MinION flow cell (*see* **Note 8**).

#### 3.3.7 Priming of the MinION flow cell and library loading

1. Thaw the tubes of Sequencing Buffer SBII, Loading beads LBII, Flush tether FLT and Flush Buffer FB at room temperature.
2. Mix by vortexing the tubes of SBII, FLT and FB, and spin down in a microcentrifuge. Keep on ice.
3. Connect the MinION Mk1B to a computer (connected to internet and having MinKNOW installed) or to a MinIT according to manufacturer’s instructions (Oxford Nanopore Technologies).
4. Open the lid of the MinION Mk1B and fit in a flow cell.
5. Open the MinKNOW software and click on ‘flow-cell check’. A MinION flow cell with more than 800 active pores is recommended for sequencing.
6. Prepare the priming mix by adding 30 µL of Flush Tether FLT into the tube of Flush Buffer FB. Mix well by vortexing.
7. Set a P1000 pipette to 200 µL and insert a 1 mL tip firmly into the priming port. Turn the wheel of the P1000 to 220-230 µL so that a small volume of yellow storage buffer is sucked up. This removes the air bubble under the cover.
8. Load 800 µL of the priming mix into the priming port, without introducing air bubbles and wait for 5 min.
9. Prepare the library for loading during the incubation time.
10. Mix the Loading beads LBII by pipetting and keep on ice.
11. In a 1.5 mL microtube, add the following reagents:

12. After 5 min, gently open the SpotON port on the flow cell.
13. Load 200 µL of the priming mix into the priming port.
14. Mix the library by tapping the tube and add into the SpotON port dropwise with a wide-mouth tip.
15. Close the SpotON port cover and the priming port.
16. Close the Mk1B lid and open the MinKNOW software to start sequencing.

|  |  |
| --- | --- |
| Sequencing Buffer SBII | 37.5 $\mu\text{L}$ |
| Loading beads LBII | 25.5 $\mu\text{L}$ |
| DNA library (20-50 fmol) | 12 $\mu\text{L}$ |
| **Total** | **75 $\mu\text{L}$** |

### 3.4 Sequencing

1. Go to ‘start sequencing’ in the MinKNOW interface.
2. Enter experiment name and sample ID. Choose flow cell type (FLO-MIN106).
3. Select kit options (SQK-LSK-110; sample type: DNA; PCR-free; multiplexing: no).
4. Select run options (Default options, *i*.*e*. length: 72 h; voltage: -180 mV).
5. To enable basecalling during sequencing, switch on the basecalling option with the ‘high-accuracy basecalling’ setting. Disable alignment during basecalling.
6. Select output location and output format (FAST5 and FASTQ).
7. Go to ‘Final review’ and then ‘Start run’ (*see* **Note 9**).

### 3.5 Data analysis

One of the advantage of Cas9-targeted sequencing is to gain access to the CG methylation (mCG) of L1 loci with high coverage. The main challenge to obtain an accurate measure of mCG levels is to discard wrongly mapped-and low-quality-reads. We describe here a user-friendly workflow which process raw reads and provides locus-specific alignment and CpG methylation levels.

#### 3.5.1 Build a custom genome

In order to distinguish the empty and filled alleles for each target (without or with the L1 insertion, respectively), we build a custom genome with contigs containing ∼55 kb per target.

1. Prepare a .bed file (`target.50kb.up.5kb.down.coordinates.bed`) in a text editor with the genomic coordinates of each target locus (starting 50 kb upstream of the L1 insertion site and ending 5 kb downstream of it), one line per locus. Note that if the relative position of the L1 element and its matched sgRNA is modified, then the intervals should be modified accordingly. For example: `chr1 95331285 95386285 ID1 0 -`
2. Prepare a similar .bed file (`L1.coordinates.bed`) in a text editor with the genomic coordinates of each target L1 (only reference elements).
3. Extract the genome sequence of 50 kb upstream and 5 kb downstream of each target L1: `bedtools getfasta -fi human.ref.genome.fa -bed target.50kb.up.5kb.down.coordinates.bed -fo target.55kb.fa`
4. For reference insertions:
  a. Prepare the file for the filled allele, which corresponds to the 55 kb sequence extracted from the reference genome: `cat target.55kb.fa > filled_allele.fa`
  b. Prepare the file for the empty allele by removing the L1 sequence: `bedtools maskfasta -fi target.55kb.fa -bed L1.coordinates.bed -fo empty_allele.fa`
5. For non-reference insertions:
  a. Prepare the file for the empty allele, which corresponds to the 55 kb sequence extracted from the reference genome: `cat target.55kb.fa > empty_allele.fa`
  b. Prepare the file for the filled allele by introducing the L1 sequence (for sense L1 use L1 sense consensus sequence, for antisense use the reverse complement sequence) at the L1 insertion point (change the chromosome number chrXXXX accordingly; for sense L1 use position=50001, for antisense use position=5001): python3 reform.py --chrom=chrXXXX --position=<50001/5001> --in_fasta=<L1.fa/L1.antisense.fa> `--ref_fasta= target.55kb.fa` > `filled_allele.fa`
6. Merge the files with empty and filled allele sequences: `cat empty_allele.fa filled_allele.fa > custom_genome.fa`

#### 3.5.2 Mapping and methylation calling

1. Index the custom genome hg38 with minimap2 and generate an index file (.mmi extension): `minimap2 -d custom_genome.mmi custom_genome.fa`
2. Create an index between fast5 and fastq files: `nanopolish index -d fast5_files ONT.fq`
3. Map ONT-fastq files to the reference genome with minimap2 and use *-a* (output alignment in the SAM format) *-x map-ont* options: `minimap2 -a -x map-ont custom_genome.fa ONT.fq > alignment.sam`
4. Sort the alignment file (SAM file) with samtools and output it as a BAM file (compressed binary version of a SAM file): `samtools view -bS alignment.sam | samtools sort -o alignment.sorted.bam`
5. Filter out reads with a mapping quality score (MAPQ) below 20: `samtools view` `-b -q 20 alignment.sorted.bam > alignment_minMAPQ20.sorted.bam`
6. Filter out reads with a mapped region shorter than 7 kb (to ensure that the mapped part of the read span the L1 sequence as well as some of the unique regions in the flanks) (*see* **Note 10**): perl -lane ‘$l = 0; $F[5] =∼ s/(\d+)[MX=DN]/$l+=$1/eg; print if $l > 7000 or /^@/’ alignment.bam > alignment_min7kmapped.bam
7. Calculate the coverage of each target. A bed file with genomic coordinates of each target is used: `bedtools coverage -a alignment_min7kmapped.bam -a target_coordinates.bed`
8. Call mCG methylation for each ONT reads: `nanopolish call-methylation –t` `4 -v -r reads.fastq -b alignment.bam -g custom_genome.fa > methylation.tsv`
9. Use the `calculated_methylation_frequency.py` script provided by nanopolish to obtain mCG level for each reference position and generate a new tab `methylation_frequency.tsv`
10. Consider only CpGs covered by at least 5 reads (*see* **Note 11**): `awk -F “\t” ‘{if($5>=5){print $0}}’ methylation_frequency.tsv > methylation_frequency_min5reads.tsv`
11. Prepare the methylation call file used to convert the bam file to a bisulfite sequencing format for the purpose of visualization in IGV: `nanomethphase methyl_call_processor -mc methylation.tsv -t 20 | sort -k1,1` `-k2,2n -k3,3n | bgzip > MethylationCall.bed.gz && tabix -p bed MethylationCall.bed.gz`
12. Convert the bam file to bisulfite bam file: `nanomethphase bam2bis –bam alignment_minMAPQ20.sorted.bam –reference custom_genome.fa – methylcallfile methylation.tsv –output OUTPUT_converted.bam`
13. Index the converted bam with samtools: `samtools index OUTPUT_converted.bam`
14. Load the converted bam file into IGV software and activate the *Color alignment by bisulfite mode* option to visualize methylated/unmethylated CpG sites (**Figure 5**).

**Figure 5.**
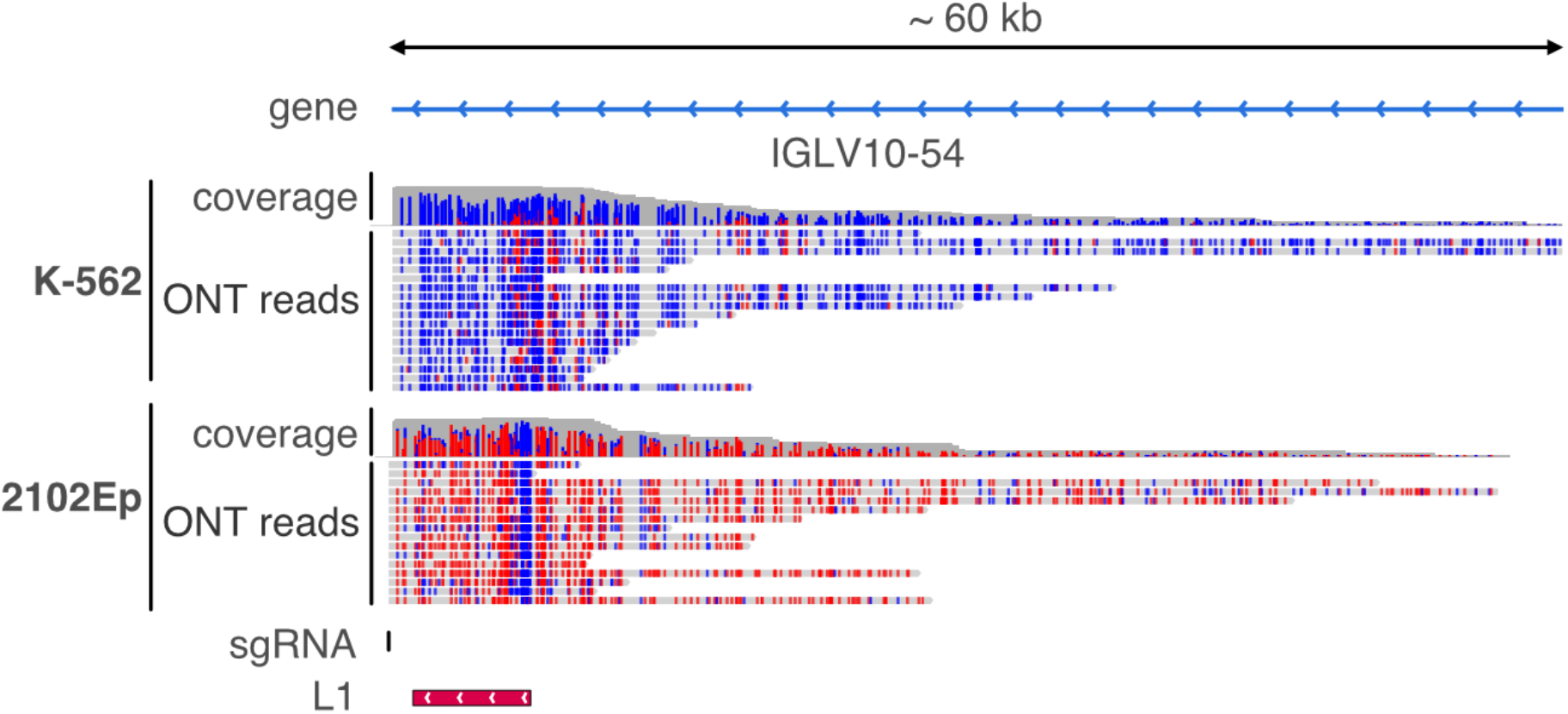
DNA methylation level of a targeted L1 locus in two different cell types. Modified IGV genome browser views of a targeted L1 locus on chromosome 22 sequenced in two different transformed cancer cells (K-562 and 2102Ep). Tracks included from top to bottom: genes; K-562 sequencing coverage and ONT reads; 2102Ep sequencing coverage and ONT reads; sgRNA position; L1 insertion position. In the coverage and ONT reads tracks, methylated and unmethylated CpG sites are highlighted in red and blue, respectively.

## 4. Notes

1. We selected sgRNAs that were pre-computed for the entire human reference genome hg38 using the CRISPOR tool [28], and made available in the *CRISPR Targets* track of the UCSC Genome Browser. However, if such a track is not available for your favorite organism, CRISPOR tool can be directly used to compute candidate targets.
2. While deciding how many sgRNAs to pool in an *in vitro* transcription reaction, remember that the greater the total number of sgRNAs, the smaller the amount of each synthesized in the pool. This amount should not be less than what is required for a successful cleavage by Cas9 at a target site. An excess of Cas9-RNP over the substrate DNA is usually recommended, with a molar ratio of 5:1 or higher [31]. In our protocol, the ratio of Cas9-RNP to DNA for each of the 125 unique target sites in the genome is approximately ∼1.5 × 10^7^:1. Therefore, we assume that this high ratio overcomes the variability in yields of individual sgRNAs during the synthesis reaction, and that it is theoretically possible to pool far more than 125 sgRNAs. Nevertheless, this remains to be experimentally tested in a MinION run.
3. We generally choose antisense sgRNA downstream of L1 to obtain the full sequence and methylation of the element and the longest possible upstream region. However, L1 can also be sequenced from its upstream flank. In this case, the sgRNA should be chosen in the sense orientation relative to L1 and upstream of it.
4. If a high number of loci are targeted for sequencing, and thus a high number of sgRNAs needs to be designed, the process should be automated to extract candidate sgRNAs from the *CRISPR Targets* Table obtained from UCSC Genome Browser, using a scripting language.
5. SgRNA LOU3161 has a unique target site in the human genome (position: chr9:83047981-83048003; strand: +). It is located approximately 1.5 kb upstream of a full-length reference L1HS insertion (chr9:83049540-83055571; strand: -) and lies within the first intron of the *RASEF* gene. It efficiently targets Cas9 and therefore can be used as an internal positive control.
6. The yield of sgRNA from the EnGen synthesis kit is 4-25 µg per reaction. Pre-warming the reagents to 37 °C and incubating for 2 h (instead of 30 min) can help improving the yield of the synthesis.
7. The original nCATS method recommends buying crRNAs (custom-designed) and tracrRNA (IDT), and assembling them individually into guide RNAs, each at 100 µM. In contrast, our approach results in lower yield per sgRNA (∼ 0.01 µM for each sgRNA in the pool, assuming equal synthesis and a yield of at least ∼4 µg of RNA after *in vitro* transcription). However, the cost for a highly multiplexed experiment is dramatically reduced and does not require aliquoting each of the assembled crRNA/tracrRNA pair.
8. We measure total DNA in the eluate rather than adapter-ligated DNA as the majority of the loaded DNA is non-ligated but may influence sequencing due to molecular crowding.
9. A caveat of nCATS is the presence of background DNA that can lead to off-target sequencing. To overcome this issue, an alternative protocol called *CABagE* [32] introduced an exonuclease digestion step to remove background DNA while genuine targets are protected by Cas9 at both ends. However, CABagE failed to reach the on-target coverage obtained by nCATS. In our experiments, background DNA does not seem to block the pores, as ∼70% of the active pores on the flow cell remain empty and are available for use throughout the run. However high molecular crowding may limit access of the target DNA to the free pores.
10. We estimate that keeping only > 7 kb mapped reads is sufficient to discard wrongly mapped reads (*i*.*e*., reads mapped solely to the body of L1). This threshold should be adapted to the length of the TE investigated.
11. We obtained sufficient coverage of each target locus to compensate for ONT sequencing errors or low-quality reads. Filtering CpG covered by at least 5 reads ensures accurate mCG level measurement over the entire L1 sequence.

## Acknowledgements

This work was supported by the French government, through the Agence Nationale de la Recherche (Idex UCAJEDI, ANR-15-IDEX-01; Labex SIGNALIFE, ANR-11-LABX-0028-01; ImpacTE, ANR-19-CE12-0032), the Fondation pour la Recherche Médicale (FRM, DEQ20180339170), Inserm (GOLD cross-cutting program on genomic variability), and CNRS (GDR 3546). We are grateful to IRCAN genomic platform, Genomed. Equipment acquisition for IRCAN platforms was supported by FEDER, Région Provence Alpes Côte d’Azur, Canceropole PACA, Conseil Départemental 06, ITMO Cancer Aviesan (plan cancer) and Inserm.

